# Inhibition of hERG K channels by verapamil at physiological temperature: Implications for the CiPA Initiative

**DOI:** 10.1101/2023.12.12.571313

**Authors:** Ashley A. Johnson, Matthew C. Trudeau

## Abstract

The Comprehensive in vitro Proarrhythmia Assay (CiPA) initiative reassesses using the inhibition of hERG channels by drugs as the major determinant for assessment of a drug’s potential to cause ‘Torsades de Pointes’ (TdP) cardiac arrhythmias. Here we report on one phase of CiPA: independent determination of hERG inhibitory properties in order to test the reproducibility of data gathered in different labs and the practicality of using the CiPA-defined recording conditions. We measured inhibition of hERG1a potassium channels stably expressed in HEK293 cells using manual whole-cell patch-clamp electrophysiology recordings at the physiological temperature of 37°C by verapamil, a known hERG inhibitor. We report that the IC50 for inhibition (180.4nM) and the Hill coefficient (1.4) for verapamil measured in an academic lab were similar to the IC50 (204nM) and Hill coefficient (0.9) measured at the Food and Drug Administration labs, indicating low variability in measurements between labs. We report that the CiPA step voltage protocol, a series of voltage steps characterized by 10 second duration depolarizing pulses at 37°C, resulted in a very low (5%) experimental success rate per cell in our hands. To circumvent the low success rate, we shortened the duration of the depolarizing pulse to 3 seconds (from 10 seconds) and shortened the duration of interpulse intervals, which is more consistent with the duration of voltage pulses used in biophysical studies of hERG, such that the entire revised protocol was shortened from over 30 minutes to approximately 10 minutes in duration. Using the revised protocol, we found an IC50 of inhibition by verapamil of 210.5 nM and Hill coefficient of 1.2. These values were similar to those generated using the original, longer CiPA step protocol. Furthermore, our success rate using the shortened protocol rose to 25%, an increase of 5-fold over the initial protocol. In summary, we captured key pharmacological data for subsequent analysis in CiPA using a revised, shorter protocol with an enhanced success rate and an overall enhanced feasibility. We propose the shorter protocol is more pragmatic for generation of hERG channel drug inhibition data for CiPA and other regulatory sciences.

## Introduction

Drug-associated, acquired cardiac arrhythmias are a common and serious clinical problem (Roden, 1993). A primary mechanism for acquired cardiac arrhythmias is the unintended, off-site inhibition of hERG potassium channels by drugs and human pharmaceuticals (Sanguinetti et al., 1995; Trudeau et al., 1995). hERG channels are the primary (alpha) subunits that form the I_Kr_ cardiac potassium channel current in native cardiomyocytes (Sanguinetti et al., 1995; Trudeau et al., 1995). I_Kr_ is a major repolarizing current in atrial ventricular myocytes (Sanguinetti and Jurkiewicz, 1990, 1991), and block of I_Kr_ current prolongs the cardiac action potential, which is a cellular precursor to arrhythmias (Sanguinetti and Jurkiewicz, 1990; Keating and Sanguinetti, 2001; Jones et al., 2014). The molecular mechanism for drug inhibition by blocking drugs is likely due to drug binding to sites in the S6 domain of the channel that lines the ion conducting pore (Mitcheson et al., 2000; Asai et al., 2021). Recognizing the link between hERG block and TdP, the FDA issued Guidance for Industry S7B and E14 which called for screening pharmaceuticals for block of hERG channels, leading to restrictions on drugs in development with the goal of promoting drug safety and minimizing cardiovascular risk (Cavero et al., 2005; FDA, 2005; FDA, 2022).

Some drugs that inhibit hERG (e.g., verapamil) were only weakly associated with pro-arrhythmia potential (Huang et al., 2017), whereas some (e.g., buprenorphine) produce QT prolongation that cannot be fully explained by hERG block (Tran et al., 2020). This indicates a potential disconnect between hERG block and QT prolongation for some drugs (Huang, 2017; Tran et al., 2020). The lack of proarrhythmic potential of some hERG inhibitory drugs could be due to their specific kinetics of block, their voltage dependence of block, or their compensatory effects on other cardiac ion channels. Thus, the potential therapeutic benefit of some drugs could be missed if a drug blocked hERG but was not pro-arrhythmic. Thus, the <u>C</u>omprehensive in vitro <u>P</u>roarrhythmia <u>A</u>ssay (CiPA) was formed to reassess the inhibition of hERG channels as the primary determinant of proarrhythmic potential and to help establish a new standard for assessing the clinical potential of drug-induced Torsades de Pointes, with the goal of enhancing drug safety without sacrificing drug usefulness (Sager et al., 2014; Fermini et al., 2016; Gintant et al., 2016; Huang et al., 2017; Tran et al., 2020; Wallis et al., 2018). CiPA has four major components: 1) to measure drug effects on human cardiac ion channels expressed heterologously in cultured cells, 2) to generate computational models of human ventricular action potentials in the presence of hERG blocking drugs using raw data gathered in 1, 3) to determine effects of hERG blocking drugs on native ionic currents and cellular electrophysiology (e.g., action potential rate and shape) in cardiomyocytes derived from human stem cells, and 4) clinical evaluation.

Here, we tested a selected reference drug (verapamil) from the CiPA initiative (Wallis et al., 2018) for inhibition of hERG channels using manual whole-cell patch-clamp at physiological recording temperature (37°C) to generate raw data as described for CiPA component 1. Verapamil is a known hERG inhibitor (Zhang et al., 1999). At room temperature (23°C), verapamil inhibited hERG channels expressed in HEK293 cells at high (143 nmol/L) affinity, likely by accessing the channel from the internal side of the membrane (Zhang et al., 1999). We also wanted to determine whether our data were similar to or different from the FDA’s in-house dataset to help evaluate data reproducibility and interlab variability using the CiPA model. Here, we report that data gathered using the CiPA step voltage protocol, which is based on depolarizing pulses of 10 sec durations (Milnes et al., 2010; Windley et al., 2017) to assess hERG inhibition by verapamil, had similar basic pharmacological properties, including an IC50 value (180.4nM), that was similar to data generated in-house by the FDA (204nM). This indicates that the conditions and protocols for hERG inhibition can be successfully reproduced at a different lab site. We found that our success rate in gathering data from cells using the CiPA step protocol was approximately 5%. To address this, we shortened the overall duration of the CiPA voltage protocol, including shortening the depolarizing voltage pulse from 10 seconds to 3 seconds, which helped us to achieve a success rate of approximately 25%. Data collected with the revised, shorter voltage protocol had an IC50 value of 210.5nM for verapamil, which was similar to the IC50 value of the longer CiPA step protocol. This result showed that the shorter voltage protocol successfully captured the basic properties of verapamil-dependent inhibition of hERG channels. We propose that the revised, shorter protocol may be more pragmatic for gathering subsequent regulatory pharmacology data using hERG channels in the CiPA initiative.

## Methods and Approach

### Cell culture and the HEK293-hERG cell line

The hERG1a-HEK293 stable cell line used here was described previously (Zhou et al., 1998) and was used for continuity with previous CiPA studies that were performed at room temperature (Windley et al., 2017). We used a cell culture handling protocol developed by Wu lab at the FDA to generate robust cells. Briefly, stable hERG1a-HEK293 cells at 75-80% confluency were washed with PBS buffer, treated with 0.25% trypsin for 30 seconds, and left undisturbed for 30 seconds after the trypsin was removed. Cells were detached with warmed DMEM solution (DMEM + 10% FBS + 10% NEAA + G-418) and gently swirled. Care was made to avoid pipetting more than necessary and to avoid introducing bubbles. Cells were either seeded at 300,000 cells in a T25 flask to passage 72 hours later or 500,000 cells to passage 48 hours later. Cells seeded at 30,000 cells per 35mM dish were plated on coverslips for manual whole-cell patch-clamp after 48 hours or 15,000 cells for recording 72 hours later to ensure single cells for whole-cell patch-clamp recordings. Compared to standard cell culture handling protocols, this protocol minimized the amount of time that the cells were handled, minimized trypsin digestion of the cells, eliminated the centrifugation and resuspension step, and established a precise amount of cells seeded. Cells that were more than 90% confluent or appeared blebby or otherwise imperfect were not used. This special handling seemed to be critical for growing cells that maintained a gigaohm seal at 37° C during the CiPA step voltage protocol. The FBS (fetal bovine serum) media supplement also appeared to be of critical importance for growing cells robust enough for the CiPA step voltage protocol. Cells grown in media supplemented with FBS that worked for our standard electrophysiology experiments (good cell growth, ability to maintain a gigaohm seal during our routine voltage protocol, and large current amplitude) would seldom withstand the CiPA step protocol. Additionally, we screened 8 lots of FBS, more than twice our standard screening procedure, to identify a suitable lot.

### Manual whole-cell patch-clamp

Whole-cell recordings were performed on hERG1a-HEK293 cell line (Zhou et al., 1998). We used an EPC10 patch-clamp and Patchmaster software (HEKA Electronic) for data acquisition. Glass microelectrodes (WPI) had 1-2 M Ohm resistances and were forged with a Sutter (P-97) electrode puller. We used an external (bath) solution containing (in mM): 130 NaCl, 10 HEPES, 5 KCl, 1 MgCl_2_*6H_2_O, 1 CaCl_2_*H_2_O, 12.5 dextrose; pH adjusted to 7.4 with 5 M NaOH; ∼280 mOsM. We used an internal (pipette) solution containing (in mM): 120 K-gluconate, 20 KCl, 10 HEPES, 5 EGTA, 1.5 MgATP; pH adjusted to 7.3 with 1 M KOH; ∼280 mOsM. These solutions had a 15mV liquid junction potential that was adjusted for in the voltage protocol. For example, -65mV was used instead of -80mV. Data was digitized at 5 kHz and filtered at 2 kHz.

Temperature control was achieved at 37°C with a Warner TC-344B temperature control device that heated the recording dish and Ringer’s solution input line to 37° Celsius. The temperature was independently verified in the bath with a thermometer. The activation of hERG channels is sensitive to temperature, becoming faster with an increase in temperature and slower with a decrease in temperature (Vandenberg et al., 2006). We found that hERG channel activation markedly sped up (the time constant of activation became smaller) when temperature was changed from room temperature (25° C) to physiological temperature (37° C). For regulatory studies, we propose that the time course of hERG activation should be reported as an independent, physiological marker for proper temperature control in the recording system.

### The CiPA step voltage protocol

The CiPA step protocol describes a sequence of voltage steps that is used to measure hERG channel inhibition by drugs for regulatory science (Windley et al., 2017; Windley et al., 2018) based around the long (10 sec) depolarizing pulses used by Milnes and colleagues to study hERG (Milnes et al., 2010). The CiPA step or Milnes protocol contains a “pulse” voltage step, which is a step to 0 mV for 10 seconds from a holding potential of -80 mV to activate outward hERG current, followed by a return step to -80 mV for 14 seconds to deactivate hERG current (Fig. 1A). The protocol also has a “no pulse” voltage step in which the voltage is held at -80 mV for 24 seconds from a holding voltage of -80 mV to keep most hERG channels in a closed state (Fig. 1B). The pulse voltage step was given 10 times, followed by the no pulse voltage step for 10 times and the sequence was repeated once, in order for the currents to stabilize (Fig. 1C,D). Stability was defined as less than 10% change in current amplitude from pulse to pulse. Drug was applied during the 2nd sequence of 10 no-pulse voltage step protocols (Fig. 1C,D), at a rate of 2 mL/min which ensured adequate solution exchange, and inhibition of hERG current was measured during the 3^rd^ sequence of pulse voltage protocols (Fig. 1C,D). The time between each pulse (or no pulse) voltage step prior to the next voltage step was 25 seconds. The holding potential of -80mV was stepped to -90 mV for 100 ms and the current response to the -90 mV step was used to calculate input resistance throughout the experiment.

**Figure 1:**
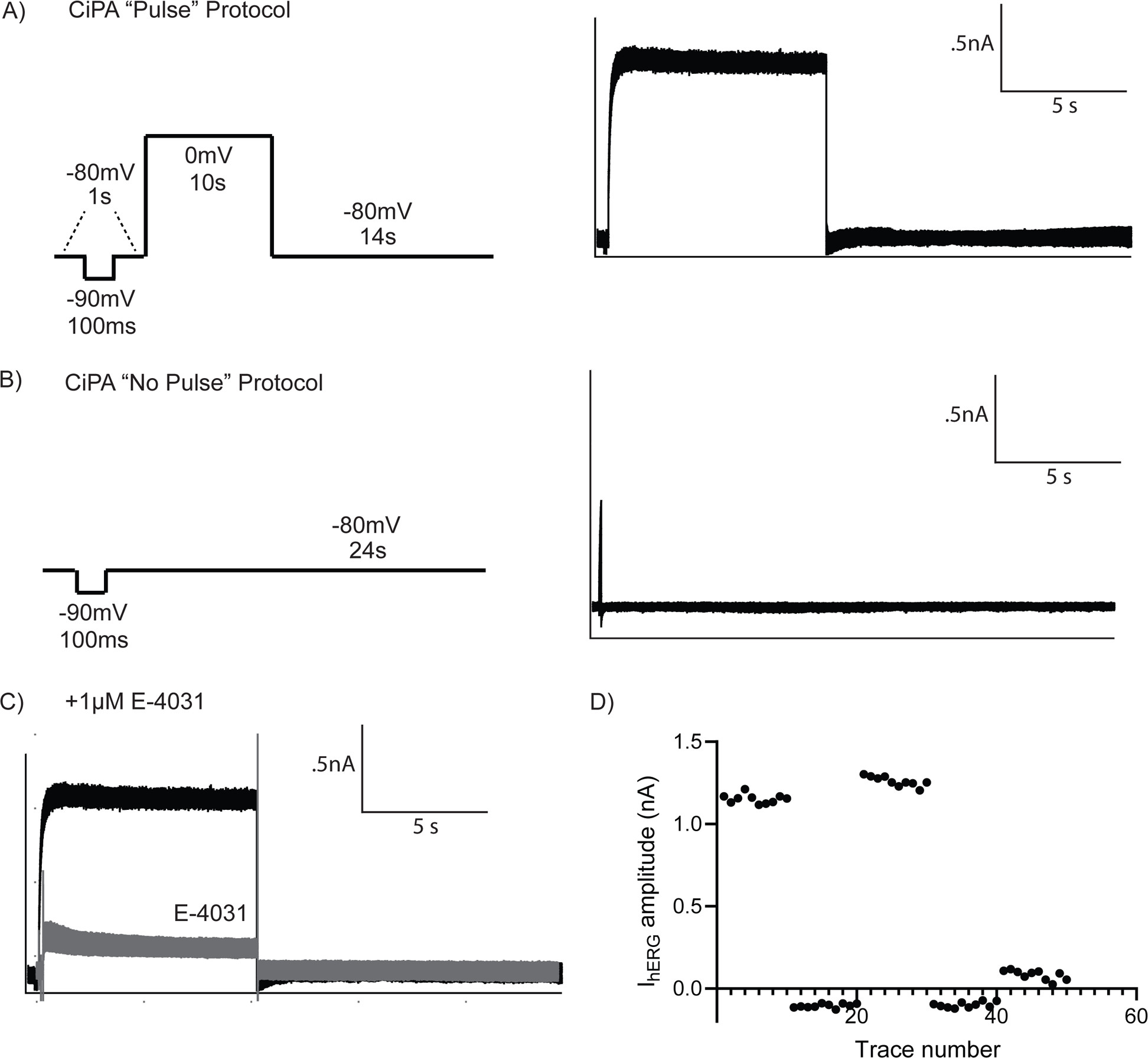
Establishing the CiPA (or ‘Milnes’) protocol at physiological temperatures. A) Left, Representation of the “pulse” voltage command featuring a 10s depolarizing voltage step to 0mV. Right, Whole-cell patch-clamp recordings of ten hERG channel currents recorded from a stable cell line (I_hERG_) measured with ten iterations of the pulse protocol depicted on the left. B) Left, Representation of “no pulse” voltage protocol as indicated and Right, Whole-cell patch-clamp recordings from the same cell as in A, but with the “no pulse” protocol. C) Representative whole-cell patch clamp current traces recorded from one cell, using the pulse protocol and with 1µM E-4031 as the test compound. D) Plot of current measured at the end of the 10 sec pulse to 0 mV or the current measured at -80 mV with the no pulse protocol versus time for a representative experiment recorded from one cell, with 1µM E-4031 applied as the test compound. Currents in C are from the final pulse protocol (black traces) and application of E-4031 (gray traces).

### Revised, shorter step voltage protocol developed in this study

We shortened the overall duration of the CiPA voltage protocol, including shortening the duration of the depolarizing pulse to 3 seconds (from 10 seconds) and shortening the duration of interpulse intervals to 9 seconds (from 25 seconds), which is more consistent with the duration of voltage pulses used in routine biophysical studies of hERG (Trudeau et al., 2011; McNally et al., 2017; Jones et al., 2018; Johnson et al., 2022), such that the entire revised protocol was shortened from over 30 minutes to approximately 10 minutes in duration. The shortened, revised protocol contains a “pulse” voltage step, which is a step to 0 mV for 3 seconds from a holding potential of - 80 mV to activate outward hERG current, followed by a return step to -80 mV for 5 seconds to deactivate hERG current (Fig. 4A). The shortened, revised protocol also has a “no-pulse” voltage step in which the voltage is held at -80 mV for 8 seconds from a holding voltage of -80 mV to keep most hERG channels in a closed state (Fig. 4B). The pulse voltage step was applied 10 times, followed by application of the no-pulse voltage step for 10 times and the sequence was repeated once, in order for the currents to stabilize (Figs. 4C,D). Stability was defined as less than 10% change in current amplitude from pulse to pulse. Drug was applied during the 2nd sequence of 10 no-pulse voltage step protocols (Fig. 4C,D), at a rate of 2 mL/min which ensured adequate solution exchange. Inhibition of hERG current was measured during the 3^rd^ sequence of pulse voltage protocols (Figs. 4C,D). The holding potential of -80mV was stepped to -90 mV for 100 ms and the current response to the -90 mV step was used to calculate input resistance throughout the experiment.

### Data Analysis

Data were analyzed off-line with Patchmaster, Igor and Prism software.

To quantify drug potency for hERG channels, the steady state hERG current amplitude from the last 1-5 hERG current traces (depending on when drug effect reaches steady-state) was compared to the amplitude from the last 5 traces measured in control solution just prior to drug application. This value is represented by one data point on the dose-response plot.

Dose response (IC50) plots were fit with the equation:

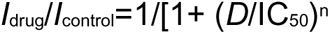

where D is the drug concentration and n is the Hill coefficient and I drug is the current in the presence of drug and I control is the current before application of drug using GraphPad Prism. Time constants of inhibition in verapamil were measured by fitting a single exponential function to the decay phase of the current in the presence of drug as in previous studies (Windley et al., 2017).

### Statistical analysis

All data are presented as mean +/- SEM. N is the number of cells. One-way ANOVA with Tukey’s test were performed with Graphpad Prism to determine statistical significance.

## Results

As a positive control, we tested the CiPA (or “Milnes”) voltage protocol (see Methods) by performing whole-cell voltage-clamp recordings at physiological temperature from a HEK293 cell line stably expressing hERG1a channels (Fig. 1). We used 10 repeats of the “pulse” protocol followed by 10 repeats of the “no pulse” protocol and repeated this sequence until hERG current was stable (Fig. 1 A, B, D). We applied the classic hERG inhibitor E-4031 (Zhou et al., 1998; Trudeau et al, 1995) at a saturating concentration (1 microM) to the bath solution during the second set of 10 repeats of the “no pulse” protocol and measured a decrease in hERG channel current during the third set of 10 repeats of the “pulse” protocol (Fig. 1C). We detected nearly complete inhibition of hERG current in the presence of E-4031 by the end of the 10 voltage pulses, as anticipated (Fig. 1D). This result indicated that the K current in the cell line was from hERG channels and that the stable cell line had no measurable endogenous K currents, which is an important consideration for regulatory studies.

We examined the inhibition of hERG by verapamil using the CiPA protocol. Verapamil is a hERG inhibitor (Zhang et al., 1999) and one of the drugs identified by the FDA as part of the CiPA panel (Sager et al., 2014; Fermini et al., 2016; Gintant et al., 2016; Huang et al., 2017; Tran et al., 2020; Wallis et al., 2018). We report that hERG currents were inhibited by increasing concentrations (30nM,100nM, 300nM and 1,000nM) of verapamil applied to the bath (Figs. 2, gray traces; 3). Verapamil inhibition was measured during the 3^rd^ series of the pulse voltage protocol following two rounds of the pulse and no pulse voltage protocols (Figs. 2, 3). We fit a curve to the dose-response plot for verapamil which yielded an IC50 of 180.4 nM (Fig. 7A). Our IC50 value (180.4 nm) was similar to the FDA’s IC50 value (204nM) for verapamil (W. Wu pers. comm) indicating good agreement between different labs (Table 1). The time constant of the hERG current decay in the presence of saturating verapamil (Fig. 2) was 4.7 (s) +/- 1.8 (s) which was similar to that of the FDA which was 10.8 (s) (95% CI of 9.6 - 12 (s)) (Fig S2, Table 1). The Hill coefficient in our data was 1.4 which was similar to that measured by the FDA which was 0.9 (95% CI of 0.7-1.1) and consistent with a stoichiometry of one verapamil molecule per channel (Table 1). Thus, the data from an independent lab (ours) was similar to the data generated in-house by a regulatory agency (the FDA), indicating low variability between labs. The IC50 values for verapamil measured here and at the FDA at physiological temperature were also similar to the IC50 (143 nmol/L) for verapamil inhibition of hERG recorded at ambient temperature (23°C) in HEK293 cells (Zhang et al., 1999), indicating that drug block of hERG by verapamil was only weakly dependent on temperature.

**Figure 2:**
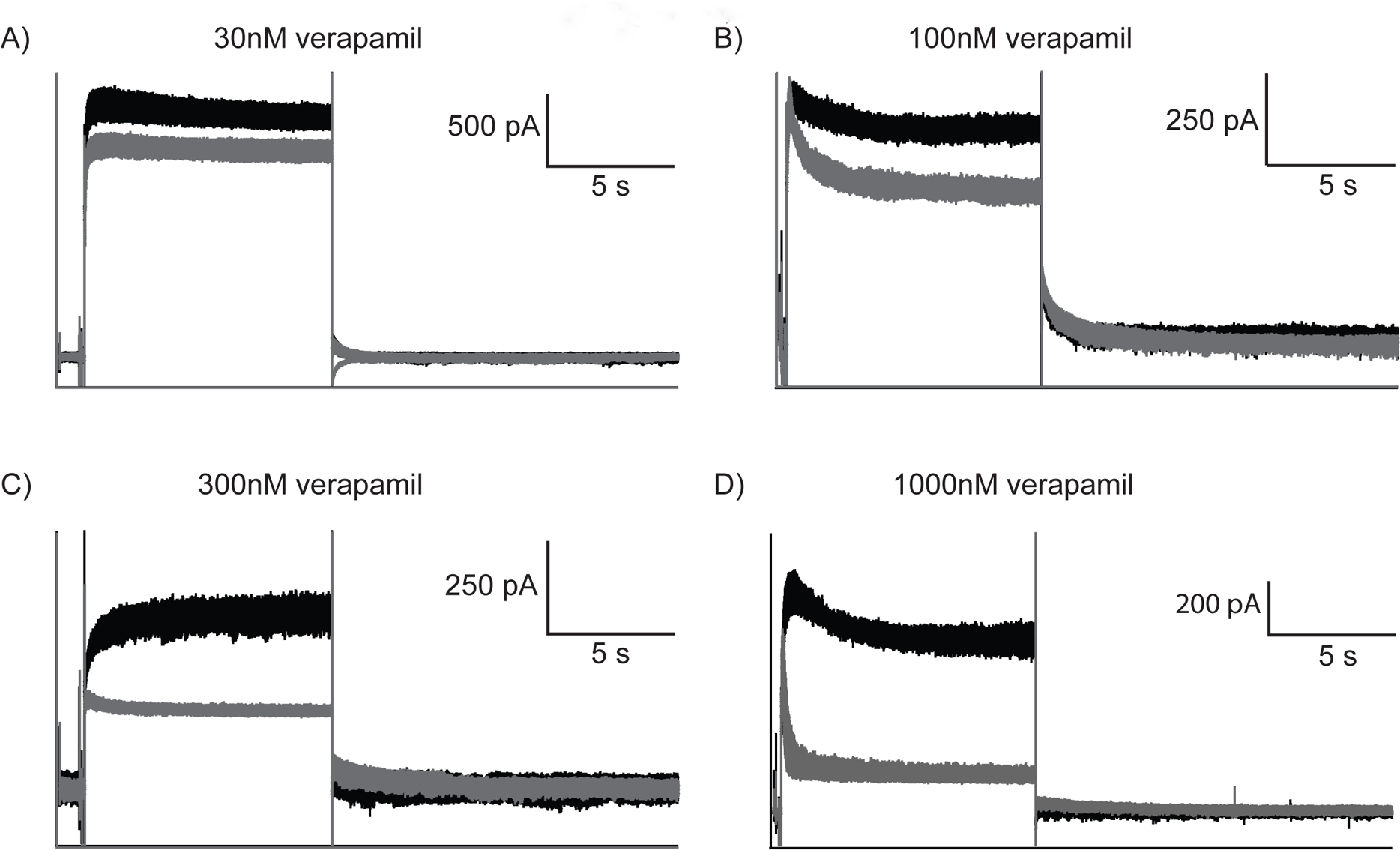
Representative current traces recorded using whole-cell patch-clamp and the CiPA voltage protocol with bath-applied verapamil. Currents before (black) and after application of verapamil (gray) at concentrations of A) 30nM, B) 100nM, C) 300nM, and D) 1000nM. Verapamil was applied during the final no pulse voltage command. Scale bars were 5 sec and the current in pA as indicated.

**Table 1:** Summary of Electrophysiology records.

|  | **Time constant**<br>**of**<br>**deactivation**<br>**(ms) at 25°C** | **Time constant**<br>**of**<br>**deactivation**<br>**(ms) at 37°C** | **IC50** | **Hill**<br>**coefficient** | **Tau of inhibition (s)\*** |
| --- | --- | --- | --- | --- | --- |
| **CiPA**<br>**Voltage**<br>**Protocol** | 399.5 +/- 20.8 | 49.6 +/- 6.3 | 180.4nM | 1.4 | 4.7 +/- 1.8 |
| **Revised**<br>**Voltage**<br>**Protocol** | 339.0 +/- 34.7 | 51.0514 +/- 3.4 | 210.5nM | 1.2 | 1.6 +/- .1 |

\*Time constant of the hERG current decay at 0mV in the presence of saturating verapamil

The success rate of our recordings using the CiPA protocol at 37° C was approximately 5%. For example, for every cell in which we achieved a gigaohm seal and began an experiment, only about 5% of cells were viable until the end of the experiment. We hypothesized that a major reason for the low success rate was the 10 second duration voltage pulse, which was a longer depolarization time than that routinely used for biophysical studies at 37°(Zhou et al., 1998; Sale et al., 2008). The 10 second duration pulses also lead to a long cumulative time (more than 30 minutes) for each experiment. With the goal of capturing similar information about verapamil inhibition of hERG but with a greater success rate, we modified the CiPA step protocol to have a shorter duration (Fig. 4). We changed the 0 mV step in the pulse voltage protocol from 10 seconds (CiPA) to 3 seconds (new) and changed the repolarization duration from 14 seconds (CiPA) to 5 seconds (new) (Fig. 4 A). We also modified the “no pulse” protocol from a duration of 25 seconds (CiPA) to 8 seconds (new) (Fig. 4B). The interpulse interval was changed from 25 seconds to 9 seconds. The number of pulses (10) remained the same. We again applied the hERG inhibitor E-4031 at a saturating concentration (1microM) to the bath solution during the second set of 10 repeats of the “no pulse” protocol and measured a decrease in hERG channel current during the third set of 10 repeats of the “pulse” protocol (Fig. 4C). We detected nearly complete inhibition of hERG current in the presence of E-4031 by the end of the 10 voltage pulses (Fig. 4D).

We used whole-cell patch-clamp to measure hERG currents using the same conditions as before (Figs. 1, 2, 3) except we applied the revised, shorter voltage protocol (Fig. 4). Verapamil was applied at the same concentration as the previous experiment (30nM,100nM, 300nM and 1,000nM) (Figs. 5, gray traces; 6) and a dose-response plot of hERG current versus verapamil concentration was fit with a curve with an IC50 (210.5 nM) which was similar to that of the IC50 value (180.4 nM) using the original CiPA step protocol (Fig. 7 B; Table 1). The time constant of inhibition in saturating verapamil at 0 mV was 1.6 (s) +/- 0.1 (s), which was similar to the value with the longer CiPA protocol which was 4.7 (s) +/- 1.8 (s) (Fig. S2, Table 1). The Hill coefficient with the new protocol was 1.2 which was similar to the original protocol. Thus, the new shorter voltage protocol successfully captured the basic properties of verapamil inhibition required for regulatory studies of hERG channels. Importantly, using the shorter protocol was also much more efficient. We used fewer cells because our success rate was five times higher, the duration of an individual experiment was shorter and the duration to generate a full data set was shorter.

**Figure 3:**
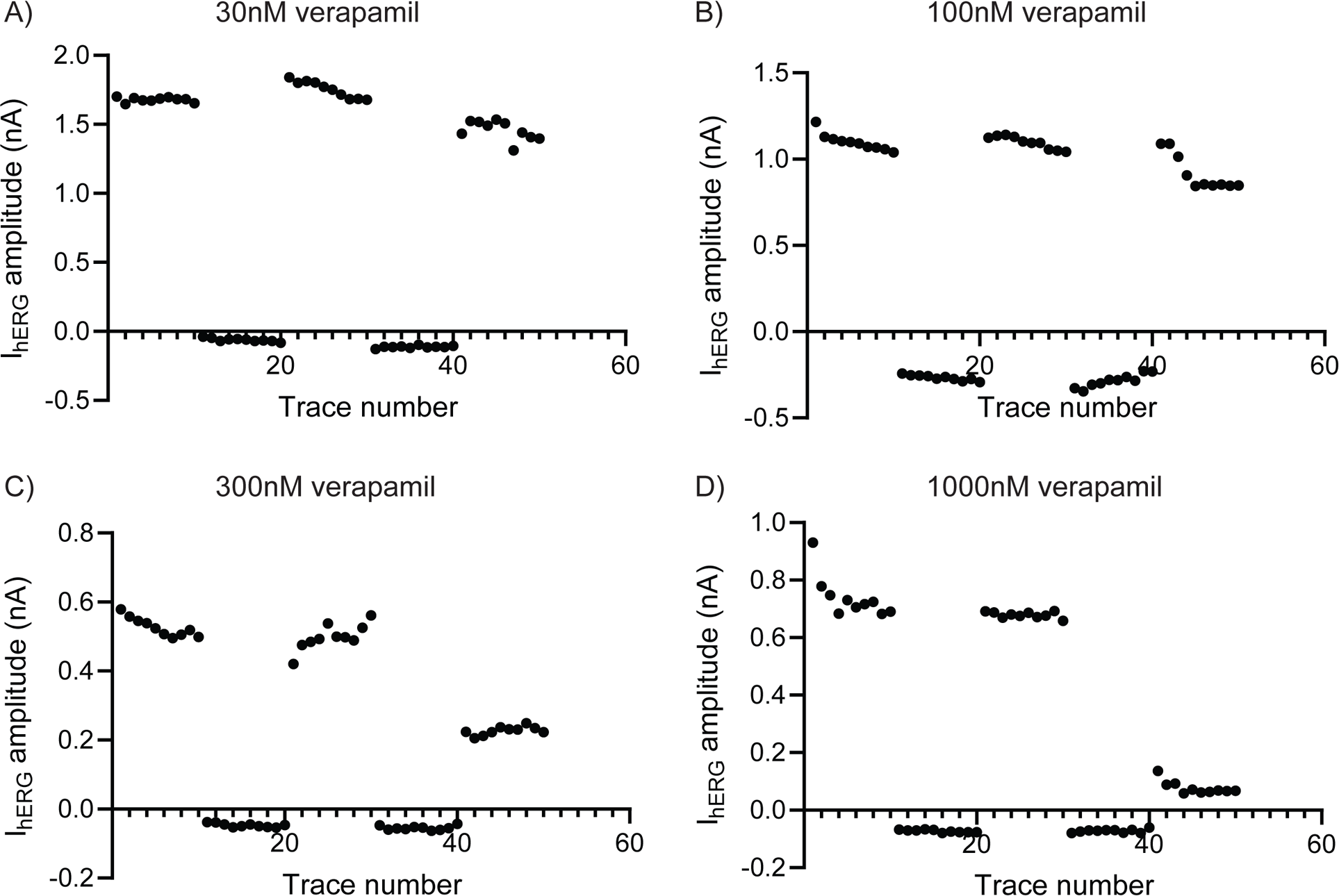
Plot of I_hERG_ peak current versus voltage trace number for a representative experiment recorded from one cell, with bath-applied verapamil at a concentration of A) 30nM, B) 100nM, C) 300nM, and D) 1000nM as the test compound.

**Figure 4:**
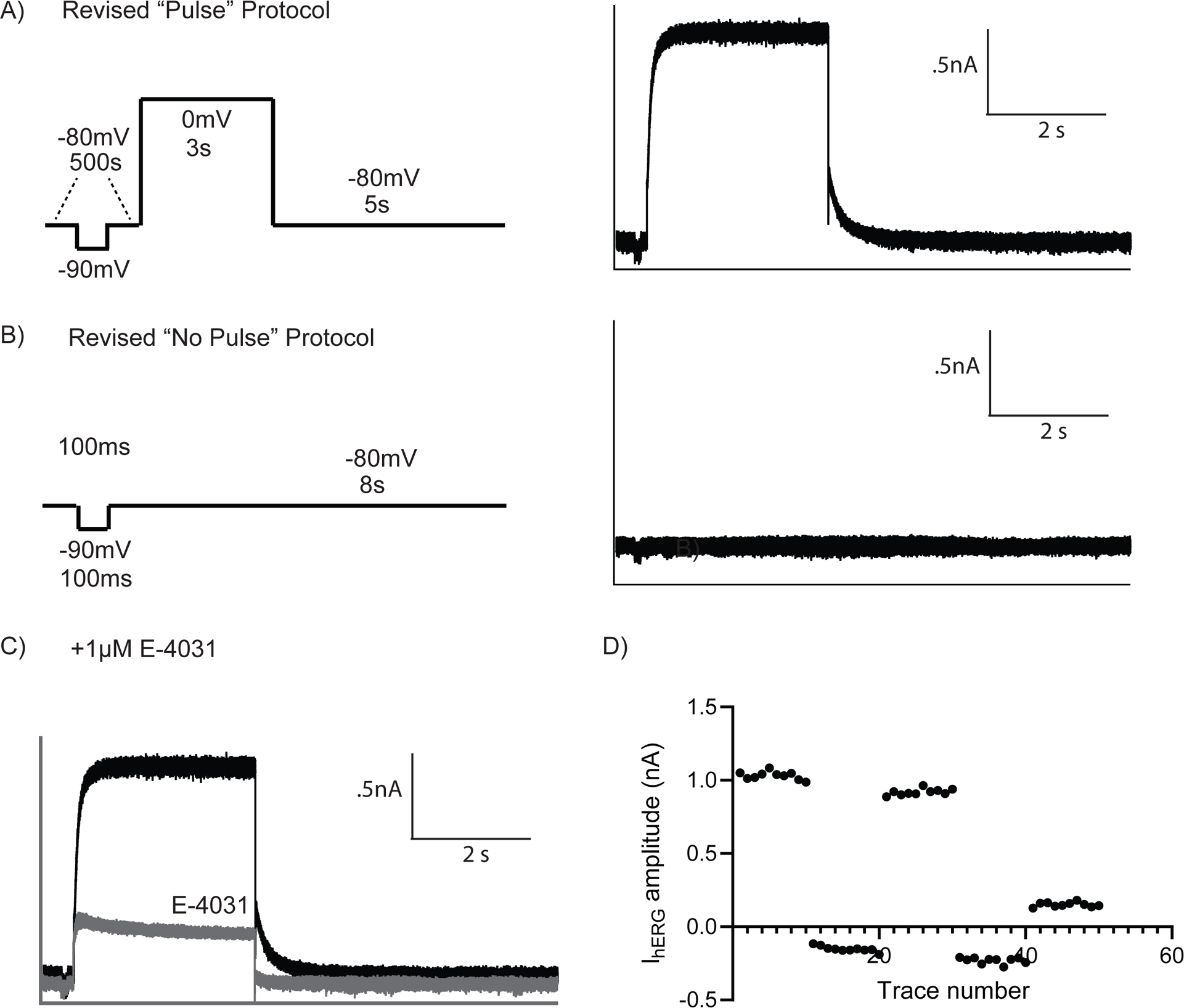
A new, revised voltage protocol with a shorter, 3-second command pulse. A), Left, Representation of the “pulse” voltage command featuring a 3s depolarizing voltage step to 0mV. Right, Whole-cell patch-clamp recordings of ten hERG channel currents recorded from a stable cell line (I_hERG_) measured with ten iterations of the pulse protocol depicted on the left B) Left, Representation of the shorter “no pulse” voltage protocol as indicated and Right) Whole-cell patch-clamp recordings from the same cell as in A, but with the “no pulse” protocol. C) Representative whole-cell patch clamp current traces recorded from one cell, using the shortened pulse protocol and with 1µM E-4031 as the test compound. D) Plot of current measured at the end of the 3 sec pulse to 0 mV (or the current measured at -80 mV) with the no pulse protocol versus time for a representative experiment recorded from one cell, with 1µM E-4031 applied as the test compound. Currents in C are from the final pulse protocol (black traces) and application of E-4031 (gray traces). Scale bars in A-C are 0.5 nA and 2sec.

**Figure 5:**
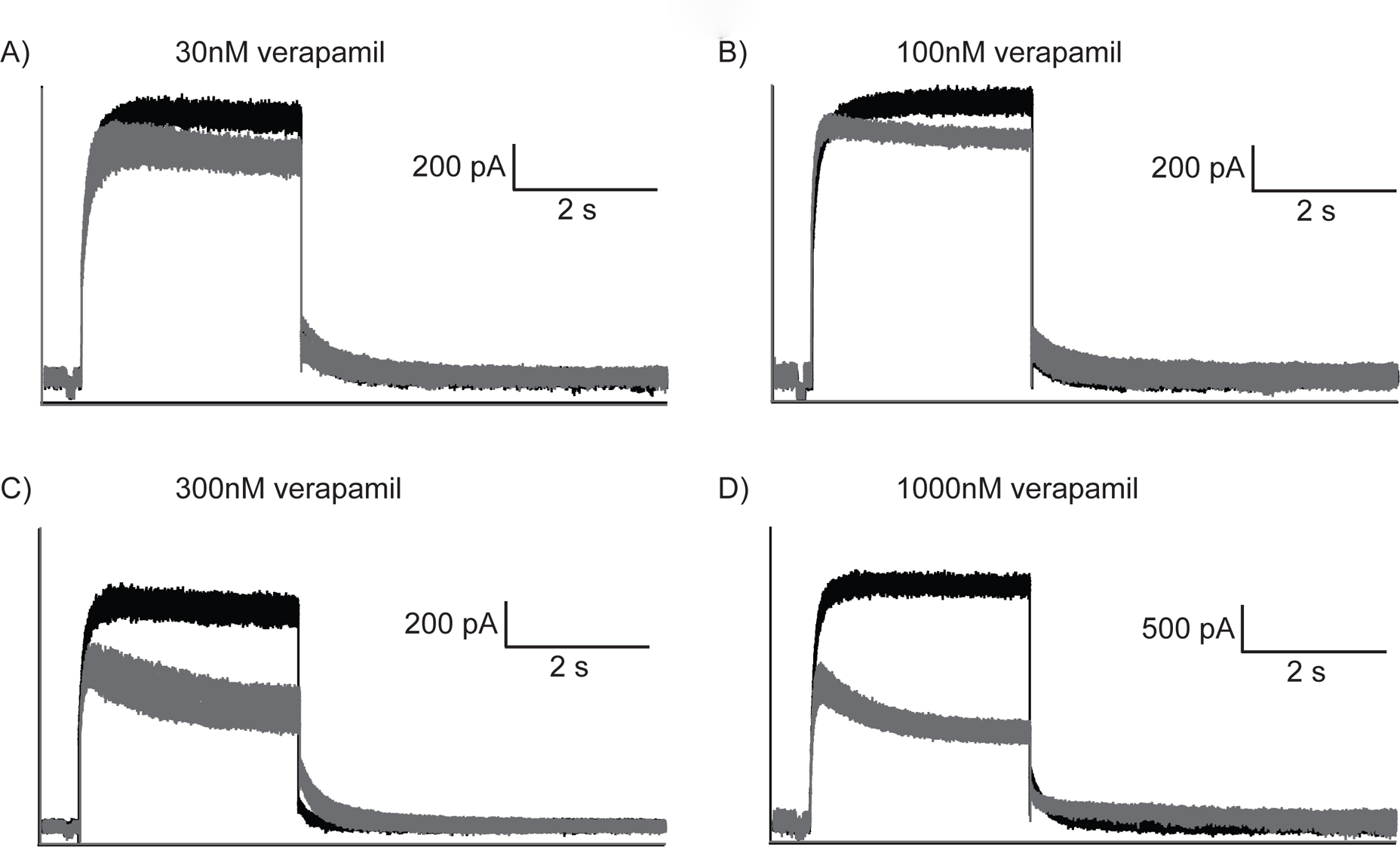
Representative whole-cell patch-clamp current traces recorded using the revised, shortened protocol indicated in Figure 4 and with verapamil to block hERG currents. hERG currents recorded before (black traces) and after application of verapamil (gray traces) at concentrations of A) 30nM, B) 100nM, C) 300nM, and D) 1000nM.

**Figure 6:**
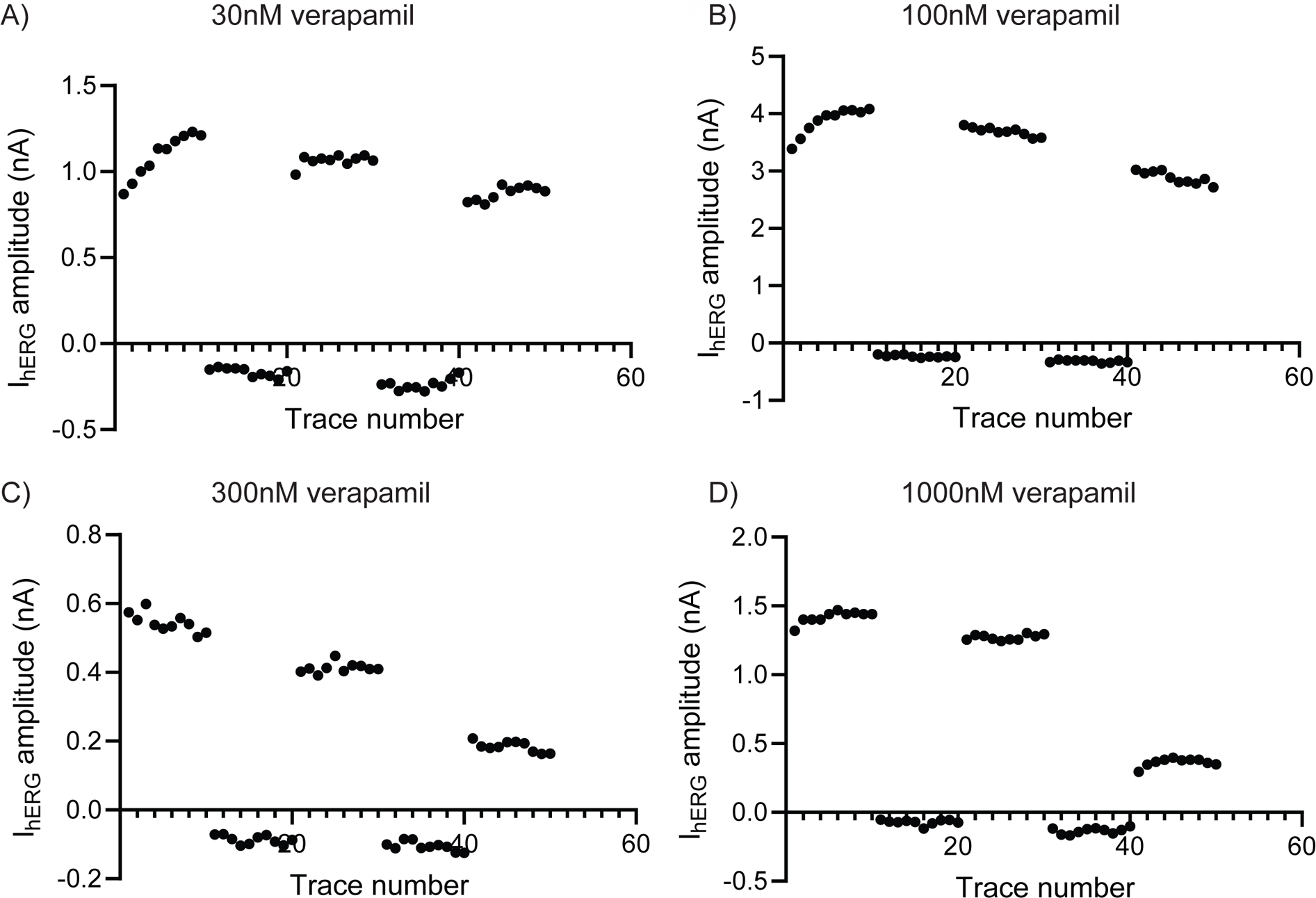
Plot of peak current amplitude versus voltage trace number for representative I_hERG_ recorded using A) 30nM, B) 100nM, C) 300nM, or D) 1000nM verapamil as the test compound.

**Figure 7:**
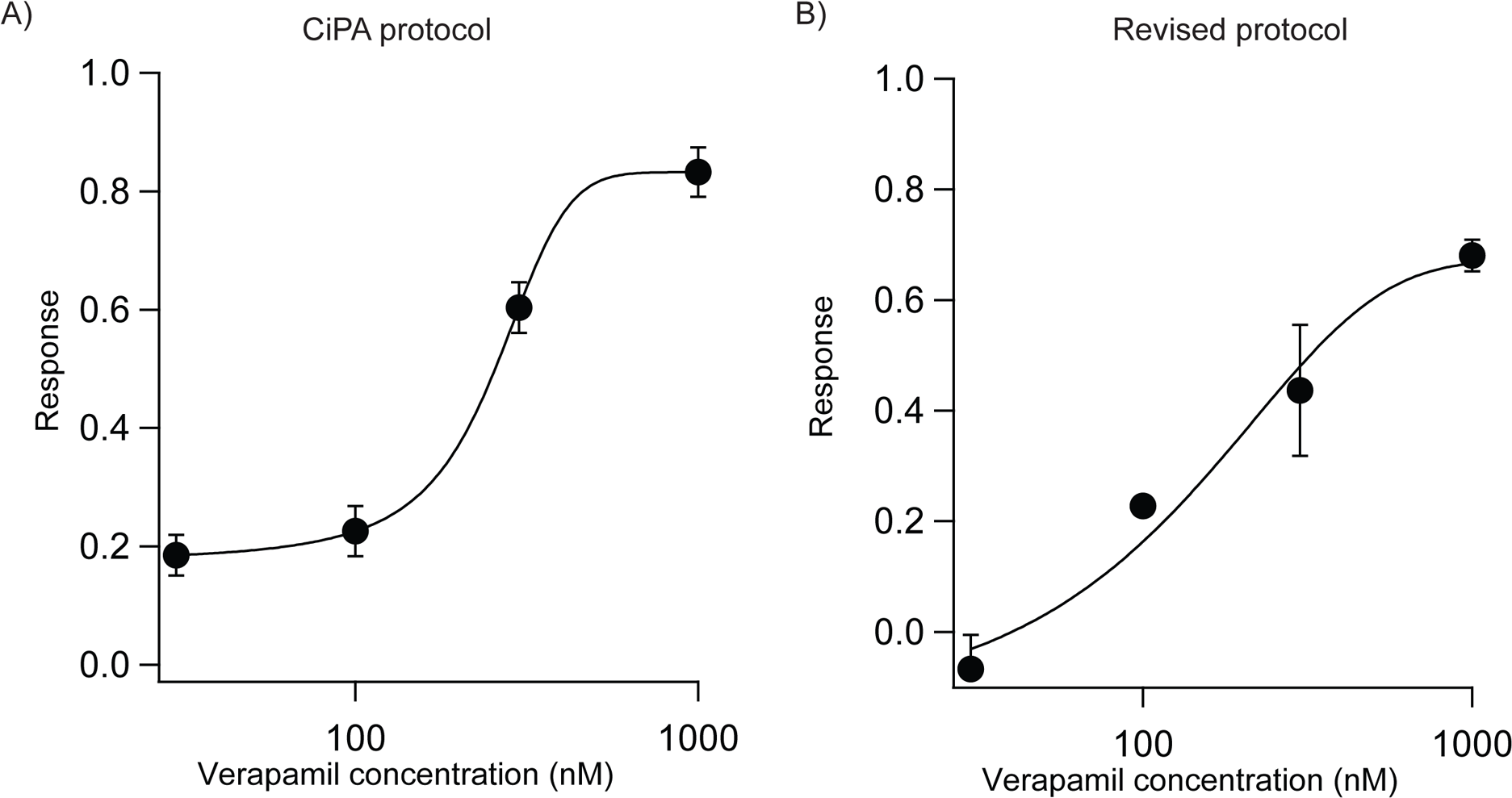
Plot of dose-response relationships. Plot of current versus verapamil concentration for A) data collected using the original 10-sec voltage step protocol and B) data collected using the revised, shorter 3-sec step voltage protocol. The IC50 determined from the original CiPA protocol was 180.4 nM and the IC50 determined using our revised, shorter (3 sec) protocol was 210.5 nM. N=3 and error bars= SEM.

## Discussion

We found that several basic pharmacological characteristics of hERG channel inhibition by verapamil, gathered in our hands, including IC50, Hill coefficient and time constant for inhibition were similar to data gathered and analyzed at the FDA. The similarities in pharmacological measurements between the two independent lab sites show that the conditions and techniques necessary to generate regulatory pharmacology data can be replicated at different lab sites, which was one of the goals of this study.

We encountered several technical difficulties in gathering hERG pharmacology data using the CiPA (or “Milnes”) voltage protocol. The gigaohm seal formed while patch-clamping would consistently fail at about the 8^th^ to 10^th^ step of the first “pulse” protocol using the CiPA protocol. This led to lack of usable data and a loss of time from the partial experiment. We estimate that approximately 5% of cells in which we formed a gigaohm seal were viable for the entire duration of the CiPA protocol. Thus, we speculate the verapamil inhibition data (Figs. 2,3) may be from a subpopulation of very robust cells. Some of these difficulties were ameliorated by using cell handling and cell culture conditions that were more stringent for regulatory sciences (see Methods) than those used for whole-cell recordings for biophysical studies. However, we suspected that most of the difficulties were due to the 10 second duration of the depolarizing pulse to 0mV, which is an extraordinarily long time to depolarize these cells and hERG channels.

We designed a new voltage protocol with a shorter depolarization pulse duration to test whether this protocol would ameliorate difficulties with the CiPA protocol. The form of the new voltage protocol was guided by the voltage protocols used routinely for biophysical and physiological studies of hERG channels to capture activation and deactivation parameters (Trudeau et al., 1995; Sanguinetti et al., 1995; Zhou et al., 1998). A 3 second depolarizing pulse to 0 mV is of sufficient duration to allow the characteristically slow hERG activation to reach steady-state, and repolarization to -50 mV for 5 seconds is of sufficient duration to measure the characteristically slow deactivation that is a hallmark of hERG. Furthermore, in our hands, the standard biophysical voltage protocols produced a high success rate for electrophysiological recordings. The revised voltage protocol had a 3 second duration for the depolarizing pulse to 0 mV and a 5 sec duration pulse for the repolarization step to -80 mV. We used a shorter duration for the no-pulse protocol, which shortened the total duration of a single voltage protocol to 8 sec. This also shortened the overall duration of the entire experiment from at least 30 min to approximately 10 min. Both the ‘pulse’ and ‘no pulse’ protocols retain a short 100ms step to -90mV to calculate input resistance throughout the experiment. Using this shorter protocol, we found that the basic pharmacological properties reported for CiPA (i.e. the IC50, Hill coefficient and time constant of inhibition) were similar for data gathered with the shorter protocol and the original CiPA protocol. Importantly, the duration of the shorter protocol was still of sufficient length for hERG currents to stabilize during two rounds of the pulse and no-pulse voltage protocols. The shorter time course was also of sufficiently long duration to measure the time course of verapamil inhibition.

We report that the revised, shorter protocol was much more efficient to use than the original, longer CiPA (or ‘“Milnes”) voltage protocol. Whereas our success rate with the original protocol was approximately 5%, our success rate with the revised, shorter protocol was approximately 20-25%. To generate the dataset with the CiPA protocol, we estimate we performed whole-cell patch-clamp experiments on at least 1,000 cells due to the low success rate of carrying out an experiment to completion. In contrast, to generate the verapamil dataset with the revised, shorter protocol, we performed patch-clamp on approximately 100 cells, due to the higher success rate of carrying out an experiment.

Based on the low success rate with the CiPA protocol we report here, we propose that it is not well justified for most regulatory pharmacology experiments. If regulatory pharmacology experiments must be performed at 37°C because, for example, hERG kinetics of activation are significantly slower at room temperature (Figure S1), then it is our recommendation that a voltage protocol consisting of 10 repeats of the “pulse” and “no pulse” protocol using a 3 second duration step to 0mV be used, as we showed here in Figure 4. hERG channel pharmacology datasets with other drugs in the CiPA panel should be gathered with this revised, shorter voltage protocol and be compared with existing FDA data sets. If the pharmacology is unchanged for other drugs using the revised, shorter protocol, then these changes will result in a higher percentage of successful experiments, a shorter time frame to complete an experiment, and thus, a shorter time to complete a collaboration or gather data for regulatory sciences. Larger data sets that do not rely on a putative subset of robust cells are likely to be more representative of the larger, general population of cells.

## References

Asai, T., N. Adachi, T. Moriya, H. Oki, T. Maru, M. Kawasaki, K. Suzuki, S. Chen, R. Ishii, K. Yonemori, S. Igaki, S. Yasuda, S. Ogasawara, T. Senda, and T. Murata. 2021. Cryo-EM Structure of K(+)-Bound hERG Channel Complexed with the Blocker Astemizole. Structure. 29:203–212 e204.

Cavero I, Crumb W. ICH S7B draft guideline on the non-clinical strategy for testing delayed cardiac repolarisation risk of drugs: a critical analysis. Expert Opin Drug Saf. 2005 May;4(3):509–30. doi: 10.1517/14740338.4.3.509. PMID: 15934857.

Fermini, B., J.C. Hancox, N. Abi-Gerges, M. Bridgland-Taylor, K.W. Chaudhary, T. Colatsky, K. Correll, W. Crumb, B. Damiano, G. Erdemli, G. Gintant, J. Imredy, J. Koerner, J. Kramer, P. Levesque, Z. Li, A. Lindqvist, C.A. Obejero-Paz, D. Rampe, K. Sawada, D.G. Strauss, and J.I. Vandenberg. 2016. A New Perspective in the Field of Cardiac Safety Testing through the Comprehensive In Vitro Proarrhythmia Assay Paradigm. J Biomol Screen. 21:1–11.

Gintant, G., P.T. Sager, and N. Stockbridge. 2016. Evolution of strategies to improve preclinical cardiac safety testing. Nat Rev Drug Discov. 15:457–471.

Huang, H., M.K. Pugsley, B. Ferminic, M.J. Curtis, J. Koernere, M. Accardia, and S. Authiera. 2017. Cardiac voltage-gated ion channels in safety pharmacology: Review of the landscape leading to the CiPA initiative. Journal of Pharmacological and Toxicological Methods. 87:11–23.

Johnson, A.C., T.R. Crawford, and M.C. Trudeau. 2022. The N-linker region of hERG1a upregulates hERG1b potassium channels. Journal of Biological Chemistry. 298(9):102233

Jones, D.K., A.C. Johnson, E.C. Roti Roti, F. Liu, R. Uelmen, A. R.A., I. Baczko, D.J. Tester, M.J. Ackerman, M.C. Trudeau, and G.A. Robertson. 2018. Localization and functional consequences of a direct interaction between TRIOBP-1 and hERG proteins in the heart. Journal of Cell Science. 131:pii: jcs206730.

Jones, D.K., F. Liu, R. Vaidyanathan, L.L. Eckhardt, M.C. Trudeau, and G.A. Robertson. 2014. hERG 1b is critical for human cardiac repolarization. Proc Natl Acad Sci U S A. 111:18073–18077.

Keating, M.T., and M.C. Sanguinetti. 2001. Molecular and cellular mechanisms of cardiac arrhythmias. Cell. 104:569–580.

McNally, B.A., Z. Pendon, and M.C. Trudeau. 2017. hERG1a and hERG1b potassium channel subunits directly interact and preferentially form heteromeric channels. Journal of Biological Chemistry. pii: jbc.M117.816488. doi: 10.1074/jbc.M117.816488.

Milnes, J.T., H.J. Witchel, J.L. Leaney, D.J. Leishman, and J.C. Hancox. 2010. Investigating dynamic protocol-dependence of hERG potassium channel inhibition at 37 °C: Cisapride versus dofetilide. Journal of Pharmacological and Toxicological Methods. 61:178–191.

Mitcheson, J.S., J. Chen, M. Lin, C. Culberson, and M.C. Sanguinetti. 2000. A structural basis for drug-induced long QT syndrome. Proceedings of the National Academy of Sciences of the United States of America. 97:12329–12333.

Roden, D.M. 1993. Torsade de pointes. Clin Cardiol. 16:683–686.

Sager, P.T., G. Gintant, J.R. Turner, S. Pettit, and N. Stockbridge. 2014. Rechanneling the cardiac proarrhythmia safety paradigm: a meeting report from the Cardiac Safety Research Consortium. Am Heart J. 167:292–300.

Sale, H., J. Wang, T.J. O’Hara, D.J. Tester, P. Phartiyal, J.Q. He, Y. Rudy, M.J. Ackerman, and G.A. Robertson. 2008. Physiological properties of hERG 1a/1b heteromeric currents and a hERG 1b-specific mutation associated with Long-QT syndrome. Circulation research. 103:e81–95.

Sanguinetti, M.C., C. Jiang, M.E. Curran, and M.T. Keating. 1995. A mechanistic link between an inherited and an acquired cardiac arrhythmia: HERG encodes the IKr potassium channel. Cell. 81:299–307.

Sanguinetti, M.C., and N.K. Jurkiewicz. 1990. Two components of cardiac delayed rectifier K+ current. Differential sensitivity to block by class III antiarrhythmic agents. The Journal of General Physiology. 96:195–215.

Sanguinetti, M.C., and N.K. Jurkiewicz. 1991. Delayed rectifier outward K+ current is composed of two currents in guinea pig atrial cells. Am J Physiol. 260:H393–399.

Tran, P.N., J. Sheng, A.L. Randolph, C.A. Baron, N. Thiebaud, M. Ren, M. Wu, L. Johannesen, D.A. Volpe, D. Patel, K. Blinova, D.G. Strauss, and W.W. Wu. 2020. Mechanisms of QT prolongation by buprenorphine cannot be explained by direct hERG channel block. PloS One. 15:e0241362.

Trudeau, M.C., J.W. Warmke, B. Ganetzky, and G.A. Robertson. 1995. HERG, a human inward rectifier in the voltage-gated potassium channel family. Science. 269:92–95.

Trudeau, M.C., L.M. Leung, E.R. Roti, and G.A. Robertson. 2011. hERG1a N-terminal eag domain-containing polypeptides regulate homomeric hERG1b and heteromeric hERG1a/hERG1b channels: a possible mechanism for long QT syndrome. The Journal of general physiology. 138:581–592.

U.S. Food and Drug Administration, HHS. International Conference on Harmonisation; guidance on E14 Clinical Evaluation of QT/QTc Interval Prolongation and Proarrhythmic Potential for Non-Antiarrhythmic Drugs; availability. Notice. Fed Regist. 2005 Oct 20;70(202):61134–5. PMID: 16237860.

U.S. Food and Drug Administration, HHS. 2022. E14 and S7B Clinical and Nonclinical Evaluation of QT/QTc Interval Prolongation and Proarrhythmic Potential--Questions and Answers: Guidance for Industry. FDA-2020-D-1791.

Vandenberg, J.I., A. Varghese, Y. Lu, J.A. Bursill, M.P. Mahaut-Smith, and C.L.-H. Huang. 2006. Temperature dependence of human ether-a-go-go-related gene K+ currents. Am J Physiol Cell Physiol. 291:165–175.

Wallis, R., C. Benson, B. Darpo, G. Gintant, Y. Kanda, K. Prasad, D.G. Strauss, and J.P. Valentin. 2018. CiPA challenges and opportunities from a non-clinical, clinical and regulatory perspectives. An overview of the safety pharmacology scientific discussion. J Pharmacol Toxicol Methods. 93:15–25.

Windley, M.J., N. Abi-Gerges, B. Fermini, J.C. Hancox, J.I. Vandenberg, and A.P. Hill. 2017. Measuring kinetics and potency of hERG block for CiPA. Journal of Pharmacological and Toxicological Methods. 87:99–107.

Windley, M.J., W. Lee, J.I. Vandenberg, and A.P. Hill. 2018. The Temperature Dependence of Kinetics Associated with Drug Block of hERG Channels Is Compound-Specific and an Important Factor for Proarrhythmic Risk Prediction. MOLECULAR PHARMACOLOGY. 94:760–769.

Zhang, S., Z. Zhou, Q. Gong, J.C. Makielski, and C.T. January. 1999. Mechanism of block and identification of the verapamil binding domain to HERG potassium channels. Circulation research. 84:989–998.

Zhou, Z., Q. Gong, B. Ye, Z. Fan, J.C. Makielski, G.A. Robertson, and C.T. January. 1998. Properties of HERG channels stably expressed in HEK 293 cells studied at physiological temperature. Biophys J. 74:230–241.

