## Supplemental Figures for "Inhibition of hERG K channels by verapamil at physiological temperature: Implications for the CiPA Initiative"

Figure S1

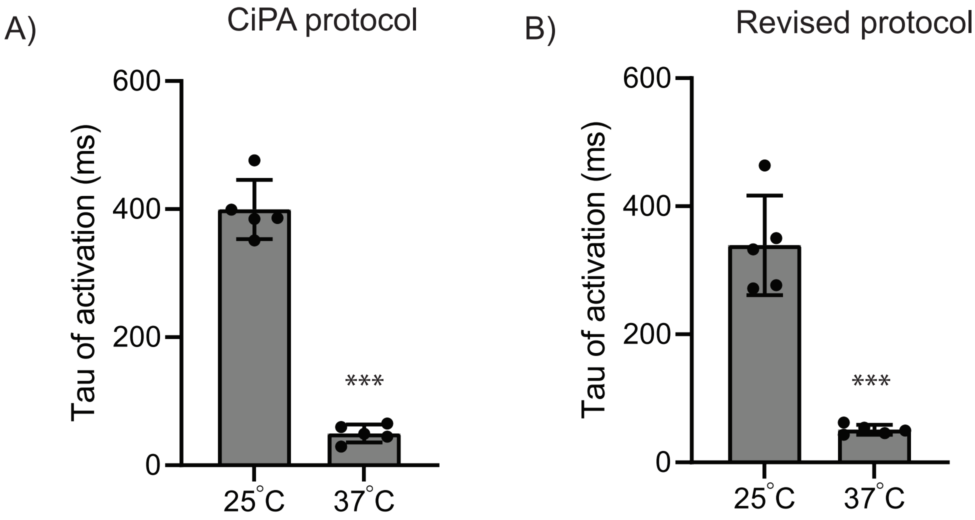
A)
CiPA protocol
600
400
200
0
Tau of activation (ms)
***
25°C
37°C
B)
Revised protocol
600
400
200
0
Tau of activation (ms)
***
25°C
37°C

Figure S2

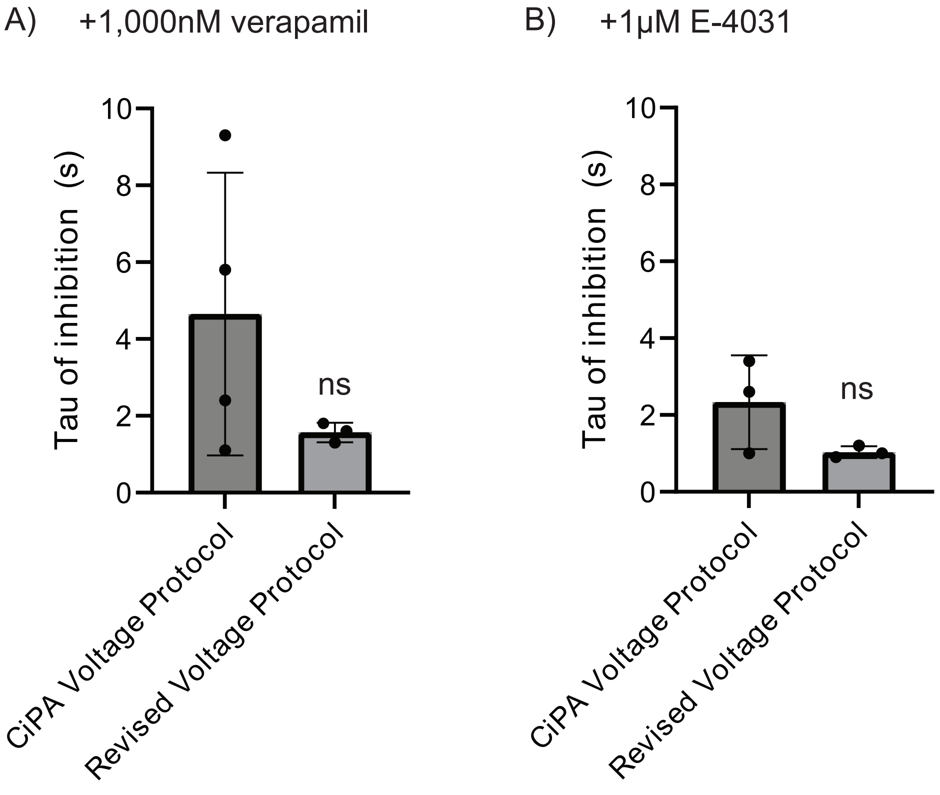
A)
+1,000nM verapamil
B)
+1µM E-4031
Tau of inhibition (s)
0
2
4
6
8
10
ns
CiPA Voltage Protocol
Revised Voltage Protocol

Figure S1: Time constant of activation for hERG channels. Plot of Tau (ms) of activation for hERG at 25°C and 37°C using A) the original CiPA protocol (data in Figures 2,3) and B) the revised, shorter protocol (data in Figures 5,6). N=4 for each.

Figure S2: Tau of inhibition for hERG channels. Plot of the time constant (seconds) of inhibition for hERG channels in A) saturating (1,000 nM) verapamil at 0mV and B) 1µM E-4031 at 0mV using both the CiPA voltage protocol and the revised, shorter protocol (indicated on figure). N=3 for each.
